# Membrane binding of pore-forming γ-hemolysin components studied at different lipid compositions

**DOI:** 10.1101/2022.02.08.479512

**Authors:** Thomas Tarenzi, Gianluca Lattanzi, Raffaello Potestio

## Abstract

Methicillin-resistant *Staphylococcus aureus* is is among those pathogens currently posing the highest threat to public health. Its host immune evasion strategy is mediated by pore-forming toxins (PFTs), among which the bicomponent γ-hemolysin is one of the most common. The complexity of the porogenesis mechanism by γ-hemolysin poses difficulties in the development of antivirulence therapies targeting PFTs from *S. aureus,* and sparse and apparently contrasting experimental data have been produced. Here, through a large set of molecular dynamics simulations at different levels of resolution, we investigate the first step of pore formation, and in particular the effect of membrane composition on the ability of *γ*-hemolysin components, LukF and Hlg2, to steadily adhere to the lipid bilayer in the absence of proteinaceous receptors. Our simulations are in agreement with experimental data of γ-hemolysin pore formation on model membranes, which are here explained on the basis of the bilayer properties. Our computational investigation suggests a possible rationale to explain experimental data on phospholipid binding to the LukF component, and to hypothesise a mechanism by which, on purely lipidic bilayers, the stable anchoring of LukF to the cell surface facilitates Hlg2 binding, through the exposure of its N-terminal region. We expect that further insights on the mechanism of transition between soluble and membrane bound-forms and on the role played by the lipid molecules will contribute to the design of antivirulence agents with enhanced efficacy against methicillin-resistant *S. aureus* infections.

**Graphical Abstract:** 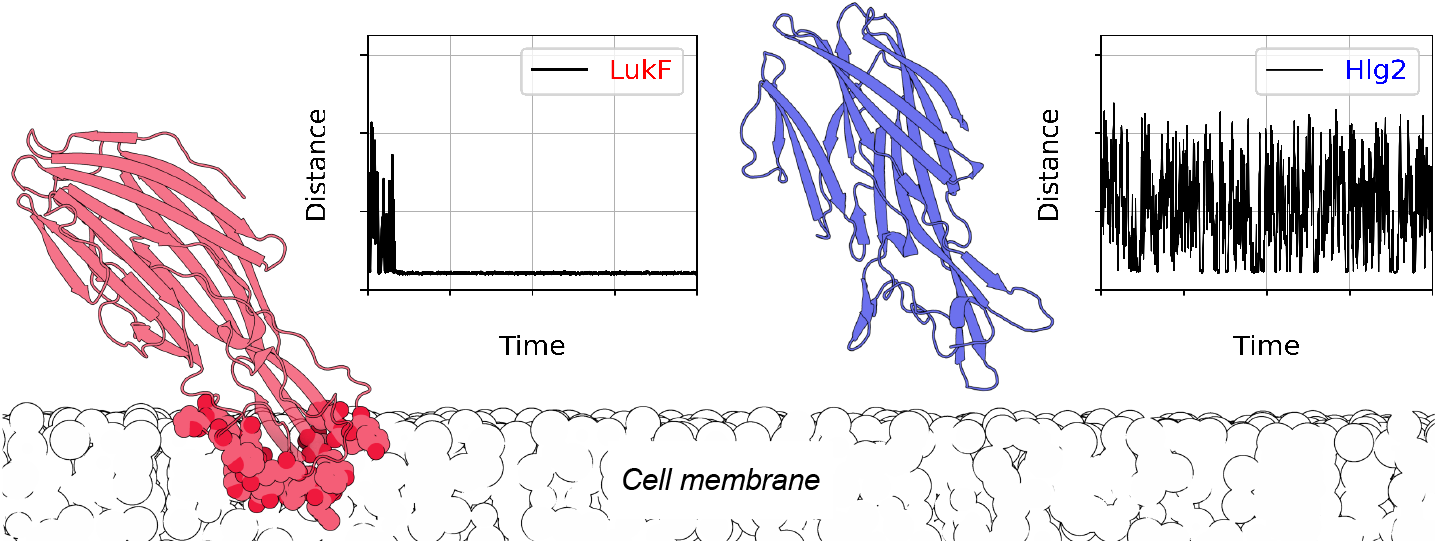

**Highlights:**

- The presence of cholesterol and unsaturated phospholipid tails facilitates the binding of *γ*-hemolysin components, LukF and Hlg2, on model membranes.
- Coarse-grained simulations show that the two components have different absorption capabilities, with LukF undergoing the most stable binding.
- The spontaneous docking of LukF on the membrane is mediated by two distant phosphatidylcholine binding sites.

## 1. Introduction

Bacterial pore-forming toxins (PFTs) are virulence factors playing a major role in bacterial pathogenesis [1]. After being secreted as soluble, monomeric proteins, they can bind to the membrane of target cells, assembling into oligomers; following a conformational change of each subunit, the PFTs penetrate the bilayer and lead to the formation of large, transmembrane pores, thus disrupting cellular homeostasis. Unfortunately, the complexity of this process poses major challenges in the study of PFTs cytotoxicity, and the detailed mechanism of pore formation is not completely understood yet [2]. Unravelling the details of the mechanism of action of PFTs would help develop ways to limit their danger, through the design of drugs that either inhibit the docking of the toxin on the cell membrane or prevent the binding between monomers. This approach would be particularly relevant for those bacterial strains showing increasing resistance towards antibiotics, as in the case of methicillin-resistant *Staphylococcus aureus.* The latter has been identified by the World Health Organization as one of the high-priority pathogens requiring urgent development of new treatments [3]; infections from *S. aureus* may indeed lead to life-threatening bacteremia, endocarditis, osteomyelitis, and necrotizing pneumonia [4].

*S. aureus* uses an arsenal of PFTs to fight the immune system of the host during infection. Among them, leukocidins are bi-component PFTs, belonging to the subfamily of *β*-barrel PFTs; their components, classified into F and S subunits, are arranged in an alternated fashion in the final pore. The octameric γ-hemolysin AB (γ-HL) is one of the most common leukocidins from *S. aureus*; the genes encoding γ-HL were indeed shown to be present in over 99.5% of human *S. aureus* isolates [5]. In addition to exerting cytotoxicity against phagocytes at the site of infection, thus providing the bacteria a way to evade host immune responses, γ-HL has been found to contribute to spontaneous pain during methicillin-resistant *S. aureus* infections, by forming pores in the neuronal membrane of dorsal root ganglia and thus directly inducing neuronal firing [6].

F and S components of γ-HL are named, respectively, LukF (also called HlgB) and Hlg2 (also called HlgA). A schematic representation of their structure in the soluble form is given in Figure 1.a. Despite the high structural similarity, the two proteins share a low sequence identity (about 30%); this difference in sequence, which is common between F and S pairs in leukocidins, accounts for the different roles attributed to the two components, with the S monomer believed to be the main determinant of cellular tropism [7]. For several leukocidins it was shown that the S components recognize and bind specific receptors on the cell surface of leukocytes; in the case of Hlg2, these receptors are CXCR1, CXCR2 and the CC-chemokine receptor 2 (CCR2) [8]. Recently, a proteinaceous receptor targeted by both LukF and Hlg2 components has also been identified, namely the atypical chemokine receptor 1 ACKR1 [9]. However, it was also experimentally observed that LukF can bind purely lipidic bilayers in the presence of phosphatidylcholine (PC) and that γ-HL possesses permeabilizing activity on model membranes in the absence of additional proteins [10, 11].

**Figure 1:**
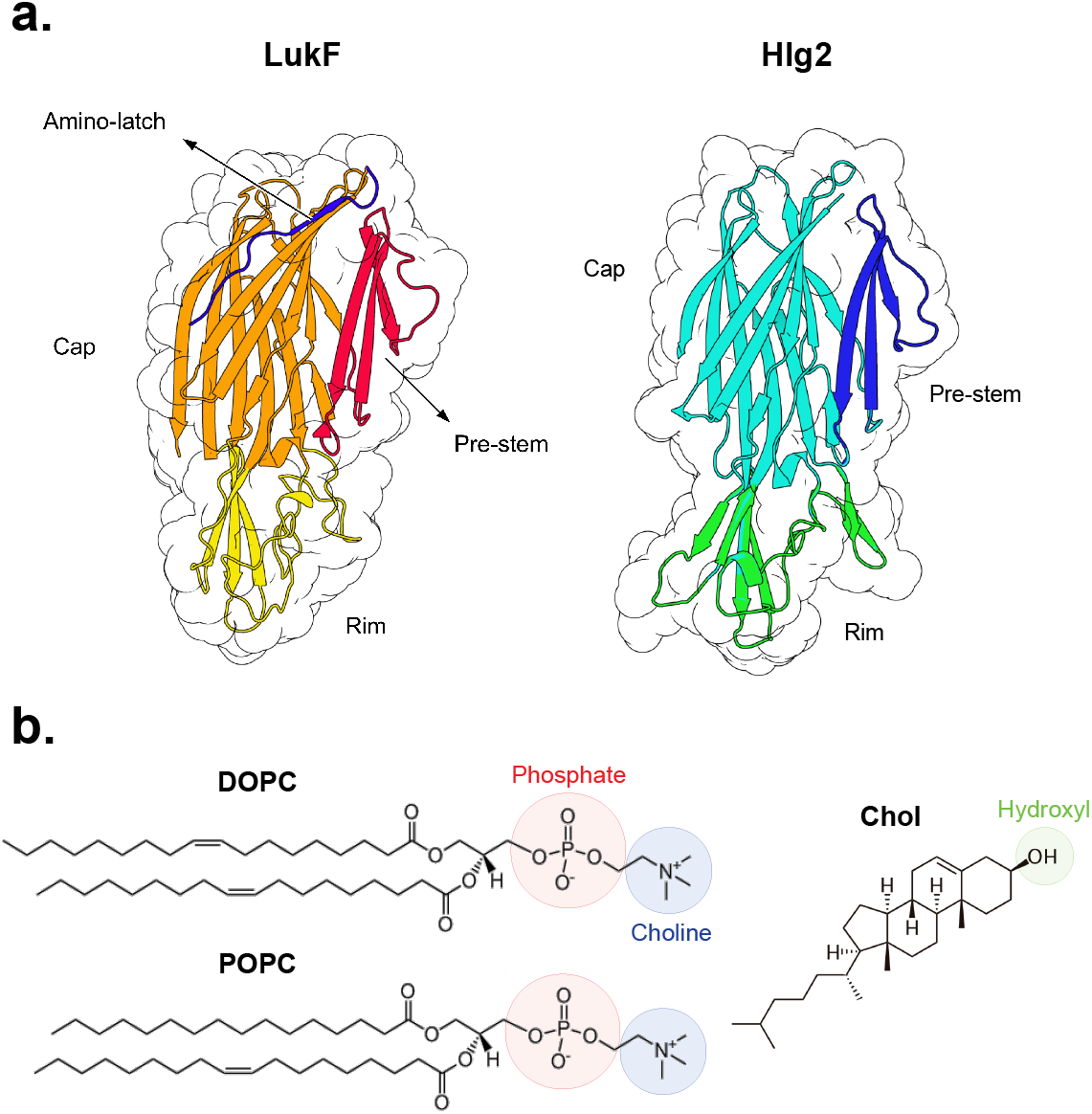
**a.** Graphical representation of the starting configurations of LukF (F component) and Hlg2 (S component) monomers. **b.** Chemical structure of the lipid molecules employed in this study.

Several residues of LukF have been previously identified as fundamental for membrane binding, through experiments performed on human erythrocyte membranes [12]; among such residues, a few have also been reported to coordinate 2-methyl-2,4-pentanediol (MPD) in the crystal structure of the fully formed γ-HL pore [13]. Arguably, there are additional lipid binding sites in LukF that have not been unveiled yet through crystallography. This is suggested by the fact that a second binding site for n-tetradecylphosphocholine (C_14_PC) has been found in recent crystal structures of other leukocidin F-components, namely LukD and LukF-PV, in monomeric form [14]. On the other hand, detailed structural information on Hlg2 docking on the bilayer membrane is currently lacking, although mutations partially disrupting some secondary structure elements of the protein, thus preventing its binding on polymorphonuclear cells, have been identified [15].

Here, we shed light on the interactions between the soluble form of γ-HL subunits and the lipid bilayer in the absence of protein receptors. To this aim, we performed a computational investigation of LukF and Hlg2 interactions with membranes at different compositions, focusing in particular on the interactions with PC components. This is attained through large-scale coarse-grained molecular dynamics simulations and state-of-the-art analysis tools of the protein-membrane interactions, which allowed us to assess the role played by the lipidic composition of the bilayer by comparison with experimental data available in literature. The experimental work by Potrich et al. [10] showed indeed a strong influence of the lipidic composition on the permeabilizing activity of γ-HL on model membranes; however, the sensitivity of each step of pore formation to the composition of the membrane could not be assessed. Through the approach presented here, we tested the hypothesis whether, and how, membrane composition affects the very first step of the perforation process, namely the protein docking on the membrane surface. We identified differences in binding caused by the presence of cholesterol (Chol) and by different degrees of saturation of the lipid tails, as in the case of 1,2-dioleoyl-sn-glycero-3-phosphocholine (DOPC) and 1-palmitoyl-2-oleoyl-sn-glycero-3-phosphocholine (POPC) (Figure 1.b); in particular, we simulated membranes of pure DOPC, DOPC:Chol (in molar ratio 1:1), and POPC:Chol (1:1). At the same time, we identify within this study the protein residues involved in direct interactions with the membrane, which are in agreement with available experimental results [12, 13].

The resulting picture sheds light on the PC-dependent membrane binding by γ-HL, highlighting the differences between the two toxin components. We expect that these results, obtained in the absence of additional membrane proteins, will help rationalize the modulation introduced by the presence of proteinaceous receptors with respect to purely lipidic systems, increase our understanding of the role played by the target membrane on leukocidins activity and facilitate the structure-based development of new antivirulence agents against *S. aureus*.

## 2. Methods

### System setup

The crystal structures of the LukF and Hlg2 components of γ-hemolysin in soluble form were obtained from the Protein Data Bank (PDB IDs 1LKF [16] and 2QK7 [17], respectively). The crystal structure of LukF lacked the disordered region between residues 130-136, which was thus modelled via MODELLER [18]. HlgB, instead, was crystallyzed as a soluble heterodimer, covalently attached to LukF through a disulfide bond; before solvation and minimization of the structure, therefore, the LukF component was removed, and C28 was mutated back to T to restore the wild type sequence. In order to study the effect of the lipid bilayer composition on the ability of the protein to anchor the membrane surface, we performed coarse-grained simulations of the monomers placed in solution at a distance of approximately 1.5 nm from the surface of the bilayer, in a simulation box of size 13 × 13 × 17 nm^3^. The MARTINI 2.2 force field was used for the whole system; in addition, the “elnedyn” representation was used for the protein to preserve its secondary structure [19, 20].

Atomistic and coarse-grained simulations of the solvated bilayers (without the presence of the protein) were performed using the force fields CHARMM36 [21] and MARTINI 2.2 respectively, in a box size of approximately 7 × 7 × 8 nm^3^. All systems were built using the CHARMM-GUI interface [22, 23], and ions were added at a 150 mM concentration. In the atomistic simulations, the TIP3P water model was employed [24]. Three membrane compositions were tested, containing: pure DOPC; DOPC and cholesterol (1:1); POPC and cholesterol (1:1). From the DOPC:Chol setup, additional binding simulations (two replicas for each protein) were performed using the MARTINI 2.3P force field [25], which includes polarizable water molecules.

### Simulation protocol

100 replicas were simulated for each membrane composition investigated. All replicas were run via the Gromacs 2019 software [26] with different initial velocities, for a total of 300 CG simulations of LukF and 300 CG simulations of Hlg2. According to the standard MARTINI procedure, a two-step minimization of the CG systems was performed, followed by a five-step equilibration phase, where the position restraints were progressively released. Production simulations were run in the isothermal-isobaric NPT ensemble; the temperature was controlled at 310 K using the velocity-rescale thermostat [27] with a coupling constant *τ_T_* = 1 ps. The pressure was controlled independently in the *xy* plane and *z* axis direction by a Parrinello–Rahman barostat [28], at a reference value of 1 bar, with a coupling constant *τ_p_* = 12 ps. Electrostatics was treated with the reaction field method, using a relative dielectric constant of 15. Electrostatic interactions were shifted to zero in the range 0 — 1.1 nm, and Lennard-Jones interactions were damped to zero in the range of 0.9 — 1.1 nm. A time step of 20 fs was used, with neighbor lists updated every 20 steps. Periodic boundary conditions were used in all directions. Each replica was run for 3 *μ*s, for an overall simulation time of 1.8 ms for the two systems. From the same starting frame used for the DOPC:Chol simulations, 2 additional 10*μ*-long simulations were performed for each protein, with the recently developed MARTINI 2.3P force field [25]. The latter, which employs a polarizable water model and a reparameterization of the cation-π interactions, was shown to reproduce with high accuracy the adsorption process of peripheral membrane proteins. This force field was here used with the aim of investigating the details of the protein-membrane interactions and the presence of lipid binding sites. In the MARTINI 2.3P simulations, the Coulombic terms were calculated using particle-mesh Ewald [29] and a real-space cutoff of 1.1 nm, and the relative dielectric constant was set to 2.5, as recommended for the polarizable MARTINI model [30]. Atomistic simulations of the bilayers were also run in the NPT ensemble; after an equilibration phase with Berendsen thermostat and barostat, in the 1*μ*s-long production runs the temperature was controlled at 310 K with the Nosé-Hoover thermostat, with *τ_T_* =1 ps; the pressure was kept at a reference value of 1 bar using the Parrinello–Rahman barostat, with a coupling constant *τ_p_* = 5 ps. PME was employed to evaluate electrostatic interactions, and a cutoff of 1.2 nm was used for all nonbonded interactions. Bonds containing hydrogen atoms were constrained using LINCS [31]. MARTINI simulations of the bilayers were run using the same procedure as for protein systems, for a duration of 1 *μs* each. A recap of the simulations performed is provided in Table S1.

### Analyses

The analyses of coarse-grained simulations were performed via Gromacs utilities [26] (specifically, *gmx mindist* and *gmx gangle*), as well as in-house python and TCL scripts.

PyLipID [32] was used to unveil the presence of lipid binding sites, and to identify the most representative binding poses. A dual cutoff was used in order to deal with the “rattling in cage” effect often encountered in CG simulations [32]; in this scheme, a protein-lipid contact is detected when their distance is smaller than the lower cutoff, and the contact is interrupted when the distance becomes larger than the upper cutoff. An analysis of the effect of the cutoffs on the detection capability of the binding sites was performed, in order to find the best values for the system under investigation. The values tested for the lower cutoff ranged from 0.425 to 0.55 nm, with steps of 0.025 nm; the values tested for the upper cutoff were 0.75, 0.8 and 0.85 nm. The simulation analysis was performed for each pair of cutoff values, and the number of identified binding sites was then plotted for each pair of cutoffs (Figure S1). Then, we followed the prescription that, when ordered according to their increasing value, the most appropriate cutoffs correspond to the largest ones right before the number of detected binding sites starts to decrease [32]. As a result, the values of 0.525 and 0.8 nm were chosen for the lower and upper cutoff, respectively.

The analyses of the atomistic and CG simulations of the bilayers were performed with Python and TCL scripts. The overall SA/lipid was computed by dividing the area of the simulation box in the xy-plane by the number of lipids per leaflet. The same quantity was also computed by taking into account the number of lipids at the level of the phosphate groups of phosphatidylcholine (average *z*-coordinate of the P atom +/− 2 Å, for each leaflet), and at the level of the choline group (average z-coordinate of the N atom +/− 2 Å, for each leaflet).

Molecular graphics of the protein have been produced with VMD [33], UCSF ChimeraX [34], and Protein Imager [35].

## 3. Results

### 3.1. LukF and Hlg2 binding is affected by phosphatidylcholine head-group exposure

The effect of the membrane composition on the binding capability of LukF and Hlg2 was investigated by considering three model bilayers, differing for the presence/absence of cholesterol (Chol) and for the degree of unsaturation of phosphatidylcholine (PC) tails: i) pure DOPC, ii) POPC and cholesterol (in ratio 1:1) and iii) DOPC and cholesterol (1:1). DOPC and POPC differ in that the former is bi-unsaturated, while the latter is mono-unsaturated. For each of the three membrane types, 100 CG simulations were performed to estimate the ability of both the F and S components of γ-HL to bind the lipid bilayer, counting the number of systems where the rim domain of the protein is in contact with the membrane at each frame (Figure 2). It is indeed generally accepted that the cell attachment of α-toxins and leukocidines’ subunits occurs by engagement of their rim domains with the headgroups of PC molecules [36, 37, 38].

**Figure 2:**
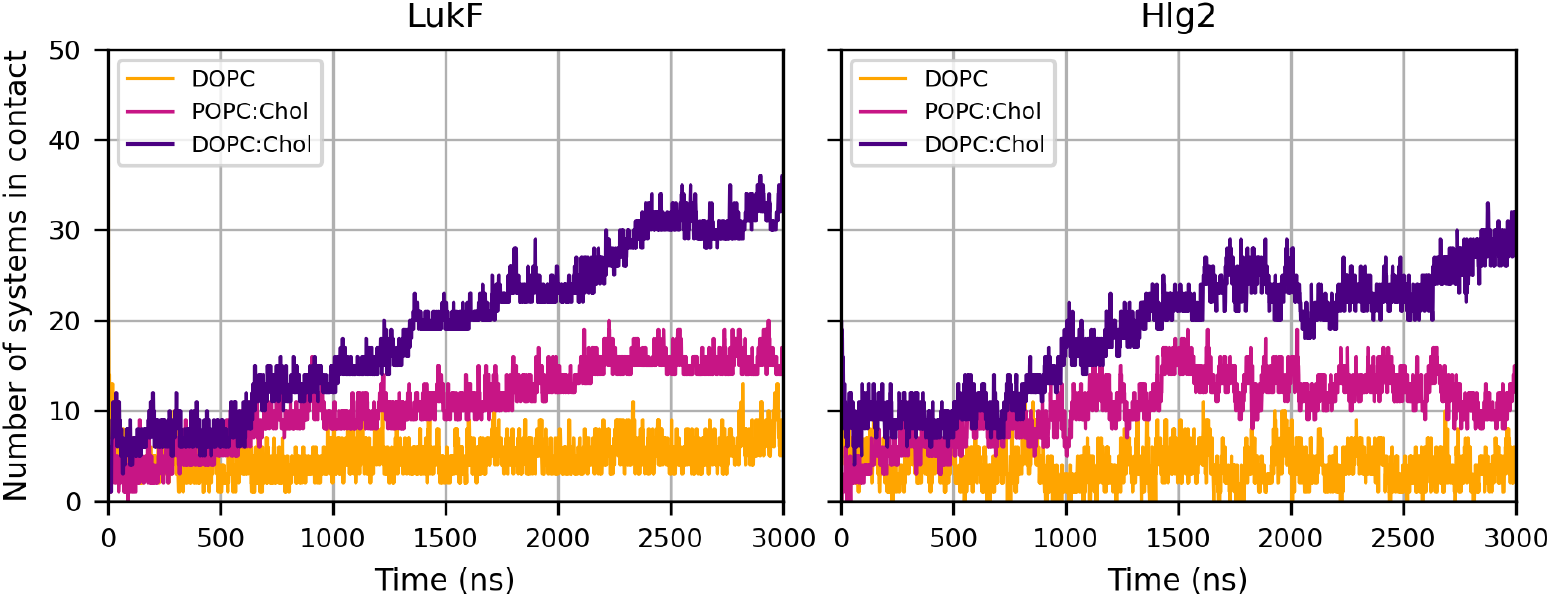
Number of systems in which the rim domain is in contact with the membrane, as a function of time. Each system was simulated in 100 independent replicas of 3 μs each.

The resulting picture from our simulations shows that monomer binding is influenced by membrane composition, thus suggesting that the latter plays indeed a key role in the very first phase of pore formation, namely protein adsorption on the bilayer. In particular, the number of bound systems follows the trend DOPC<POPC:Chol<DOPC:Chol. These results are in line with experimental data suggesting a dependence of pore formation on membrane composition [10]: in this study the formation of pores in pure lipid model membranes was quantified by monitoring the permeation of calcein with fluorescence spectroscopy. The average percentage of calcein release measured in the experiments was 0 for the DOPC membrane, 45.3 ± 3.0 for POPC:Chol (1:1), and 67.7 ± 7.7 for DOPC:Chol (1:1). Although we cannot rule out the effect of the membrane on other steps of pore formation, such as oligomerization and folding of the β-barrel, the qualitative agreement between the experimental results and our simulations suggests that the effect of membrane composition in the absorbtion step is a key determinant of the overall toxin efficiency.

In search of a molecular rationale for these results, we compared computed observable quantities from both atomistic and CG simulations of the solvated bilayers, reproducing the same lipidic compositions employed before (Table 1). In particular, we focused on the accessibility of the headgroups of PC molecules to the protein by measuring the surface area per lipid (SA/lipid); this was suggested to be deeply affected by the chemical composition of the membrane on the basis of the so-called “unbrella model” [39, 40, 41]. The latter accounts for the fact that cholesterol, given the small size of the polar, hydroxyl group, tends to be buried under the large polar heads of PC, thus limiting water exposure; however, if the lipid chains of PC are short or unsaturated, cholesterol can hardly fit below the PC headgroups, thus intercalating and further separating the lipid molecules.

**Table 1:** Surface area per lipid (SA/lipid) computed from the atomistic simulations of the hydrated bilayers. SA/lipid is computed also at the level of the phosphate group and the quaternary ammonium cation of the choline group (P and N, respectively).

| Resolution | Composition | SA/lipid [nm<sup>2</sup>] | SA/lipid (P) [nm<sup>2</sup>] | SA/lipid (N) [nm<sup>2</sup>] |
| --- | --- | --- | --- | --- |
| Atomistic | DOPC | $0.68 \pm 0.01$ | $0.70 \pm 0.02$ | $0.73 \pm 0.03$ |
| | POPC:Chol | $0.42 \pm 0.01$ | $0.79 \pm 0.03$ | $0.86 \pm 0.03$ |
| | DOPC:Chol | $0.45 \pm 0.01$ | $0.79 \pm 0.04$ | $0.90 \pm 0.03$ |
| Coarse-grained | DOPC | $0.68 \pm 0.01$ | $0.77 \pm 0.03$ | $0.88 \pm 0.04$ |
| | POPC:Chol | $0.43 \pm 0.03$ | $0.88 \pm 0.03$ | $1.06 \pm 0.05$ |
| | DOPC:Chol | $0.45 \pm 0.01$ | $0.93 \pm 0.04$ | $1.13 \pm 0.05$ |

To quantify lipid accessibility, we first computed the overall SA/lipid (first column in Table 1) by taking into account the total number of lipids per leaflet. The results for atomistic and CG simulations are very similar, thus confirming the accuracy of the MARTINI model. In addition, these data are in agreement with experimental values of SA/lipid available in literature, obtained by SAXS or NMR experiments (Table S2). Comparing the values obtained for the different membrane compositions, the average SA/lipid is large for DOPC, but low and very similar for the other two cases. This reflects the well-known fact that cholesterol molecules are tightly packed with the lipid tails: given its small polar group, cholesterol tends to be buried in the hydrophobic layer of cholesterol-rich membranes, partially under the large polar heads of phospholipids, with restricted orientation fluctuations [42]. This phenomenon tends to increase the overall SA/lipid. In addition, the value for POPC:Chol is slightly smaller than the DOPC:Chol, due to the saturation of one of the lipid tails of POPC with respect to DOPC, thus leading to a more ordered and efficient packing of the hydrophobic portion of the molecules.

On the contrary, the SA/lipid computed at the level of phosphate groups (second column in Table 1) is slightly larger for the cholesterol-containing membranes than in the pure DOPC case, thus suggesting a higher accessibility of the polar heads when cholesterol is present. Indeed, both *in vitro* and *in silico* experiments have shown that the addition of cholesterol to fluid bilayers causes significant structural changes, including increased headgroup spacing [43] and hydration [44] (in addition to increased bilayer thickness [45, 46] and reduced water penetration into the membrane hydrocarbon region [44, 47]). Moreover, the SA/lipid computed at the average z-coordinate of the nitrogen atom of the choline groups increases from POPC:Chol to DOPC:Chol. Given the recognized importance of the formation of cation-π interactions between PC and aromatic residues of peripheral proteins, including bacterial toxins [48, 49], the choline exposure is expected to be very important for the anchoring of the protein and the stabilization of its docked pose, thus explaining the large number of binding events observed between the protein and the DOPC:Chol membrane.

The structural features of the bilayers are therefore able to appropriately describe the extremely low number of binding events of LukF and Hlg2 to pure DOPC membranes, where the absence of cholesterol causes a packing of the lipid heads. In addition, this analysis clearly explains the effect determined by the degree of unsaturation in DOPC:Chol and POPC:Chol membranes: unsaturated acyl chains increase the spacing among PC headgroups, thus improving the possibility for the protein residues to form contacts. In the next section we will investigate in greater detail how the headgroup exposure caused by the presence of cholesterol and bi-unsaturated lipid chains affects the docking of LukF and Hlg2, and which residues are involved in this protein-membrane interaction.

### 3.2. Similarities and differences in the membrane binding of LukF and Hlg2

As observed in the previous section, the trend followed by the absorption capability on different membranes is similar for both LukF and Hlg2 (Figure 2). However, in the first case the number of systems with successful binding tends to increase over time, or to fluctuate around a constant value when a plateau is reached; in the case of Hlg2, instead, larger fluctuations are observed. These are clear indications of a lower stability of Hlg2 binding with respect to LukF.

In line with this argument, Figure 3.a displays the average residence time of the toxin on the bilayer, computed as the average duration (over 100 replicas) of the longest uninterrupted contact between the protein and the membrane. This quantity increases from DOPC, to POPC:Chol, to DOPC:Chol, following the same trend of the number of contacts; in addition, the residence time is slightly lower for Hlg2 with respect to LukF for all membrane types under investigation. These results are consistent with the average number of binding/unbinding events reported in Figure 3.b: indeed, the number of unbinding events is larger for the pure DOPC membrane. This can be explained by recalling that, given the low accessibility of choline groups of PC in such system, a stable binding can not be easily attained; at the same time, the high density of such groups on the surface of the cholesterol-lacking membrane might facilitate electrostatic interactions with the protein. The overall effect is a large number of binding/unbinding events, with a short residence time. In addition, we observe that in the cholesterol-containing membranes the number of unbinding events is larger for Hlg2 than for LukF, consistently with the larger fluctuations in the number of bound systems reported in Figure 2.

**Figure 3:**
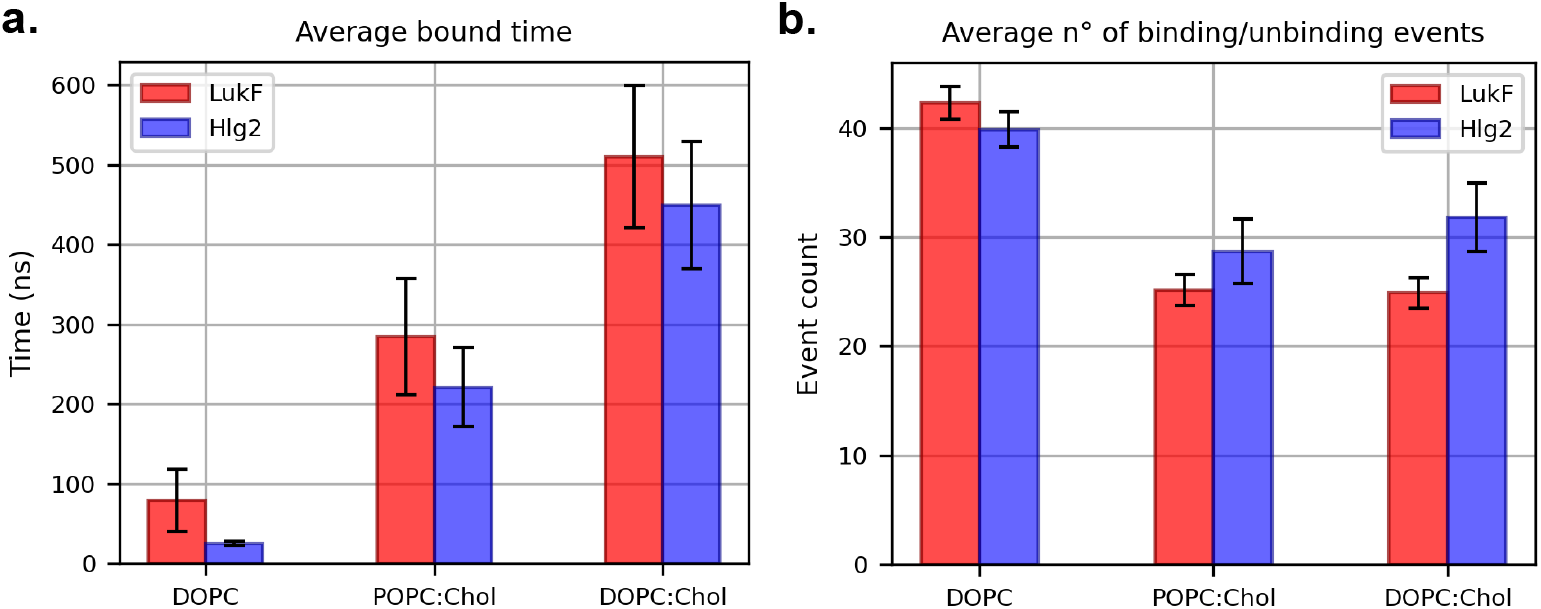
**a.** Average residence time of the protein on the membrane surface, at different membrane compositions. **b.** Average number of binding/unbinding events between the rim and the membrane in the course of a simulation.

We performed electrostatic calculations on the crystal structures of LukF and Hlg2, aimed at understanding how electrostatics drives the interaction of each monomer with the membrane (Figure 4). Both proteins show a largely positive electrostatic potential (LukF bears a +7 net charge, while Hlg2 a + 14 net charge). As expected, a comparison between the two monomers shows the complementarity of the areas forming inter-protomer interfaces in the assembled pore, namely sides I-II and sides II-I of LukF and Hlg2, respectively. In addition, an unexpected difference between the two proteins is observed in the rim domain: Hlg2 lacks the deep, negatively charged pocket found on the bottom of the LukF rim, which is likely to facilitate and further stabilize the contact between the protein and the positively charged choline groups of PC.

**Figure 4:**
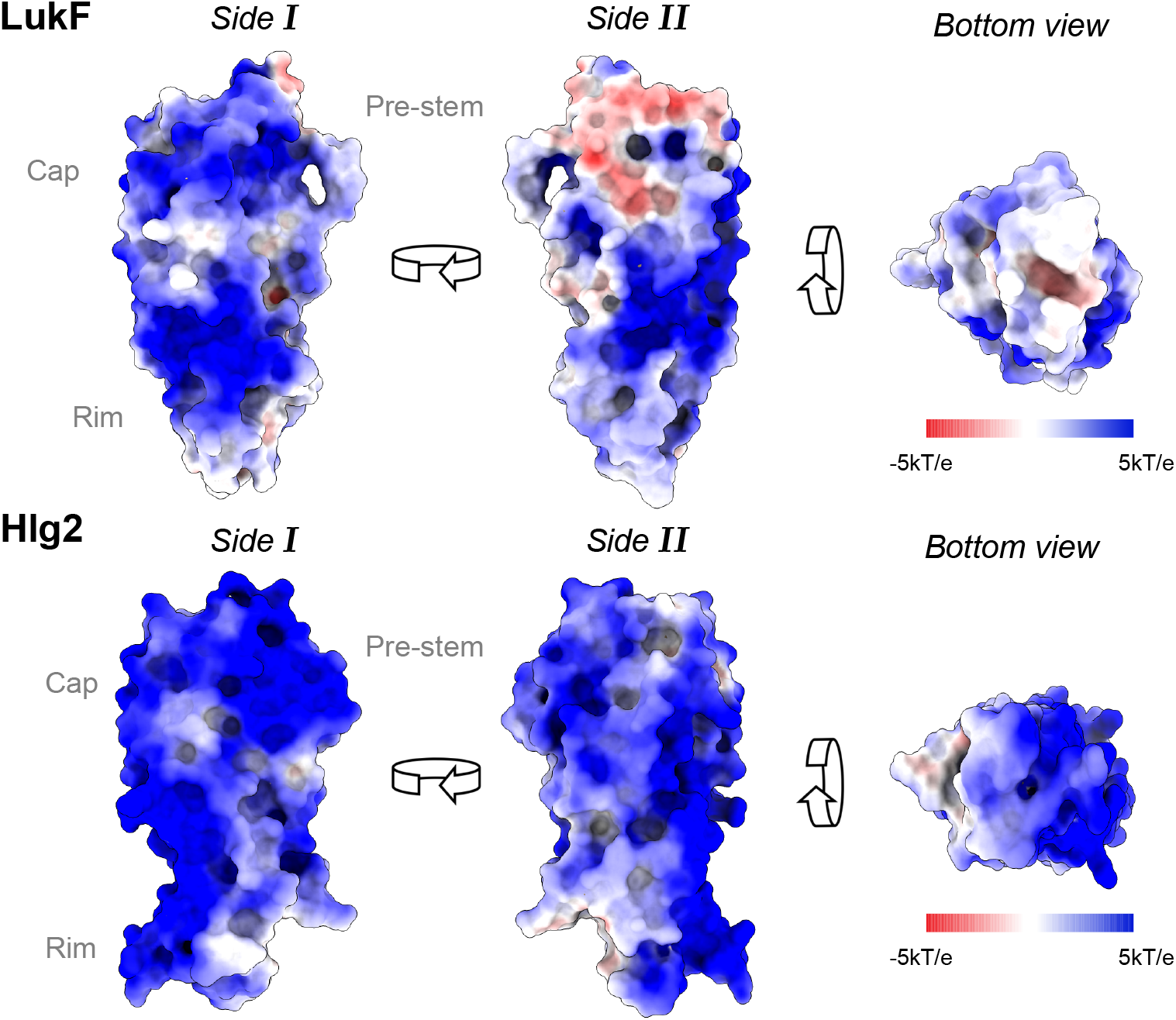
Electrostatic potential calculated with the adaptive Poisson-Boltzmann solver (APBS) [50], mapped on the surface of LukF and Hlg2 monomers.

A more detailed insight is provided by the per-residue number of protein-membrane contacts (Figure 5.a,b). For both proteins, there are substantial changes in the profile of the residues interacting with the membrane at different compositions. These changes are associated with a different orientation of the protein on the membrane, as quantified by the distributions of values of the angle formed between the z-axis and the axis along the anchored protein (Figure 5.c; for the definition of the protein axes, see Figure S3).

**Figure 5:**
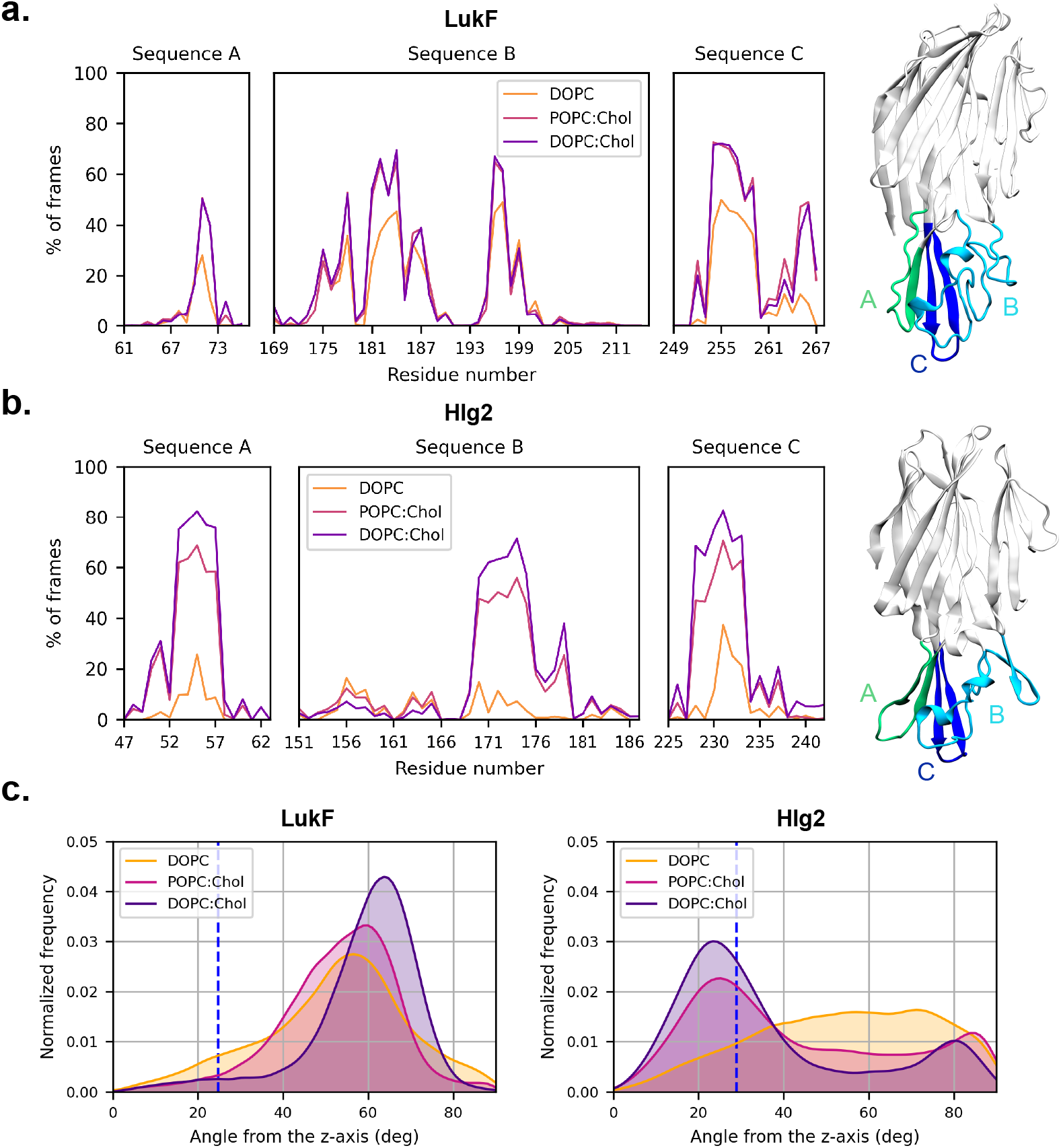
Percentage of frames in which each residue is involved in a contact with the membrane for the rims of **a**. LukF and **b**. Hlg2. The three distinct sequences forming the rim domain are also coloured on the protein structures for clarity. Data for the full protein sequence are plotted in Figure S2. **c.** Distributions of the angles formed between the protein axis (as defined in Figure S3) and the axis perpendicular to the membrane. The dashed lines correspond to the angle formed by the protomers in the crystal structure of the assembled pore (PDB ID: 3B07).

In Hlg2, the inability to efficiently bind PC in the pure DOPC membrane leads to an unspecific binding, with a broad distribution of protein orientations and the lack of rim residues forming a predominantly large number of contacts. Increasing the exposure of PC heads (as in POPC:Chol and DOPC:Chol), more selective interactions are established between the segments A (residues 48 to 64) and C (residues 225 to 242) of the rim and the membrane; at the same time, the involvement of the first segment of region B (turn 155-157, distant in space from the other loops of the rim region) is reduced, while the remaining part of the same segment (residues 158 to 188) is involved in a larger number of interactions. This is associated to a change in the orientation of the protein, with its main axis slightly aligning to the normal to the bilayer.

We notice that Hlg2 predominant orientations, although very close to the one adopted by the protomer in the structure of the assembled pore [13], lead to a less stable binding with respect to LukF, where we observe the opposite trend. In fact, the positioning of LukF on the membrane surface is different from the final position of the protomer in the assembled pore, where the main axis of the protein is almost perpendicular to the membrane plane. The single monomer, instead, is bent towards the membrane for those lipidic compositions corresponding to an increased affinity.

The profile of the number of contacts in LukF is very similar for POPC:Chol and DOPC:Chol (Figure 5); however, in the second case, the larger number of contacts involving residues Q254 and Y267, both located on the upper region of the segment labelled C, determines a slight bending of side II of the monomer towards the membrane with respect to the POPC:Chol case, thus explaining the slight shift in the angle distribution. In addition, the increasingly bent orientation of LukF toward the membrane plane in the cases of higher affinity is very specific, leaving only one dimerization interface exposed toward the solvent (side I, namely the one that is “locked” by the N-terminal, the so-called amino-latch). This asymmetry emerges from Figure S2, where we observe for LukF a large number of contacts with the membrane in the sequences 150-160 and 290-299, both belonging to side II. This specific orientation of LukF on the membrane might be facilitated by the asymmetric charge distribution on the two faces of the rim domain, with side II characterized by a more extended surface with a less positive electrostatic potential (Figure 4). Moreover, the bent orientation is stabilized by some polar/charged residues located on a loop of the cap domain, on the opposite side with respect to the amino-latch. These residues (R156, N159, Y160, K161) interact specifically with the polar heads of PC in the cholesterol-containing membranes.

The observations above are confirmed by additional simulations of LukF and Hlg2 monomers binding on DOPC:Chol bilayers, performed with the MARTINI 2.3P force field. LukF steadily binds to the membrane after approximately 900 ns, and remains bound for the rest of the 10 μs-long simulations. Hlg2, on the other hand, interacts with the membrane for short time intervals, never achieving a proper stable binding (Figure 6.a). The resulting picture is even more dramatic than the one emerging from Figure 3, and is in line with previous experimental observations that reported the binding of Hlg2 on DOPC:Chol liposomes as almost completely reversible, at variance with the more stable binding of the LukF component [11]. In addition, it was shown that, in some conditions, LukF can attach to the membrane surface independently on the S subunit and that the primary binding of LukF on the membrane is necessary for Hlg2 binding, as shown on the study of the pore-forming activity of γ-HL on human erythrocytes [51]. This picture, however, may change in the presence of proteinaceous receptors of Hlg2 [7]; in such cases, the primary binding of Hlg2 might promote the secondary interaction between LukF and the cell surface, as observed in studies of γ-HL activity on human polymorphonuclear cells [15].

**Figure 6:**
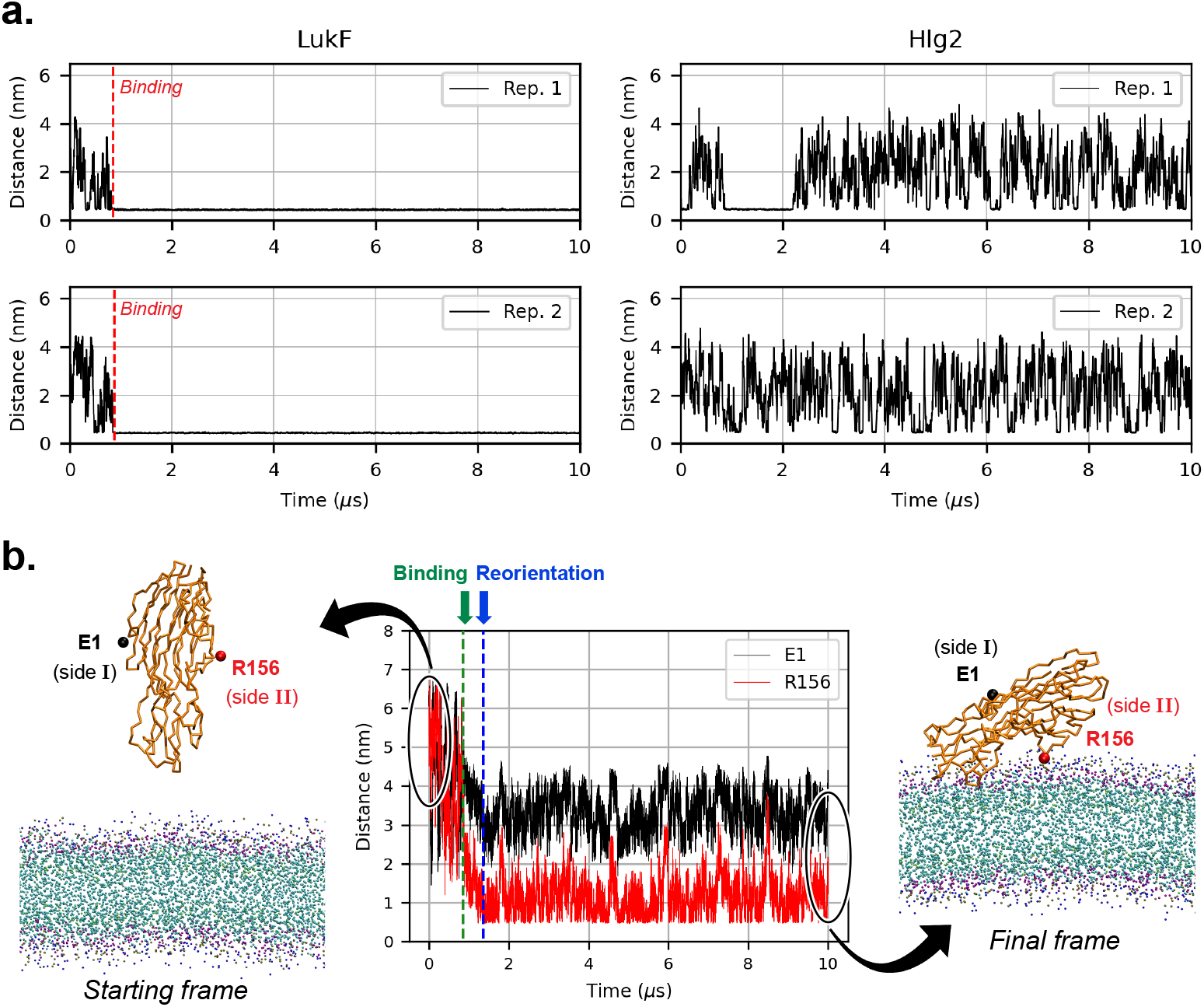
**a.** Minimum distance between the protein and the membrane in the two replicas performed with the MARTINI 2.3P force field. **b.** The interaction between LukF and the membrane is mediated through the side II of the monomer, as shown by the minimum distance between the bilayer and the backbone beads of E1 (side I) and of R156 (side II). The membrane composition is here DOPC:Chol, and the system is simulated with the MARTINI 2.3P force field (replica 1).

Our additional CG simulations confirm also the asymmetric orientation of LukF. The evolution of the protein orientation in both replicas is here monitored by measuring the minimum distance between two residues located on two opposite faces of the monomer, namely E1 on side I and R156 on side II (Figure 6.b and Figure S4). In replica 1, docking takes place in two steps: first membrane binding, followed by protein reorientation. In replica 2 the second step does not take place, since the toxin binds to the membrane in an orientation compatible with the stable pose of replica 1; this is demonstrated by the per-residue distances between the protein and the membrane in Figure S5, which displays a comparison between the binding/reorientation events of replica 1 and the binding event of replica 2. We suggest that the specific orientation of LukF on the membrane might facilitate the interaction with Hlg2 components diffusing in solution, through the exposure of a dimerization interface towards the solvent. In this picture, Hlg2, although unable to steadily bind to the lipid bilayer, could anchor the membrane surface by binding the exposed interface on the F component; the successive interaction of Hlg2 with its proteinaceous receptor or with a second hete-ordimer would then drive the reorientation of the complex on the membrane, aligning the protein axes closer to those of the protomers in the octamer. A similar hypothesis was formulated in [13] on the basis of the position of the membrane-binding residues in LukF crystal structure; however, the interface supposedly exposed to the solvent was the opposite with respect to the one revealed by molecular dynamics simulations of the binding event. In our picture, a key role for dimerization seems to be played by the region surrounding the N-terminal, which locks the dimerization interface in the soluble form of LukF; therefore, unbinding of the N-terminal is expected upon interaction with the incoming Hlg2 monomer. In the next sections we will describe how these results provide also new insights into the mutagenesis data of lipid binding.

### 3.3. Details of LukF/membrane interactions

The simulations of membrane binding performed with the polarizable MARTINI 2.3P force field were used to analyze at a higher detail the interactions between the toxin monomers and the membrane, using the recently developed python package PyLipID [32]. Given the lack of a stable binding pose for Hlg2, we focus our analysis on the LukF monomer. Moreover, we consider only the case of the DOPC:Chol membrane, given its particularly high propensity to promote LukF binding with respect to other lipidic compositions investigated.

In particular, we identified two main binding sites on the LukF monomer (Figure 7). Both of them presented a 100% occupancy, defined as the percentage of frames in which the lipid-protein residue contacts are formed. The first binding site (BS1) involves residues Y72, D73, F74, T188, R256, W257, N258, G259, F260; the second one (BS2) includes residues W177, G178, P179, Y180, H186, Y189, E192, R198, Y261.

**Figure 7:**
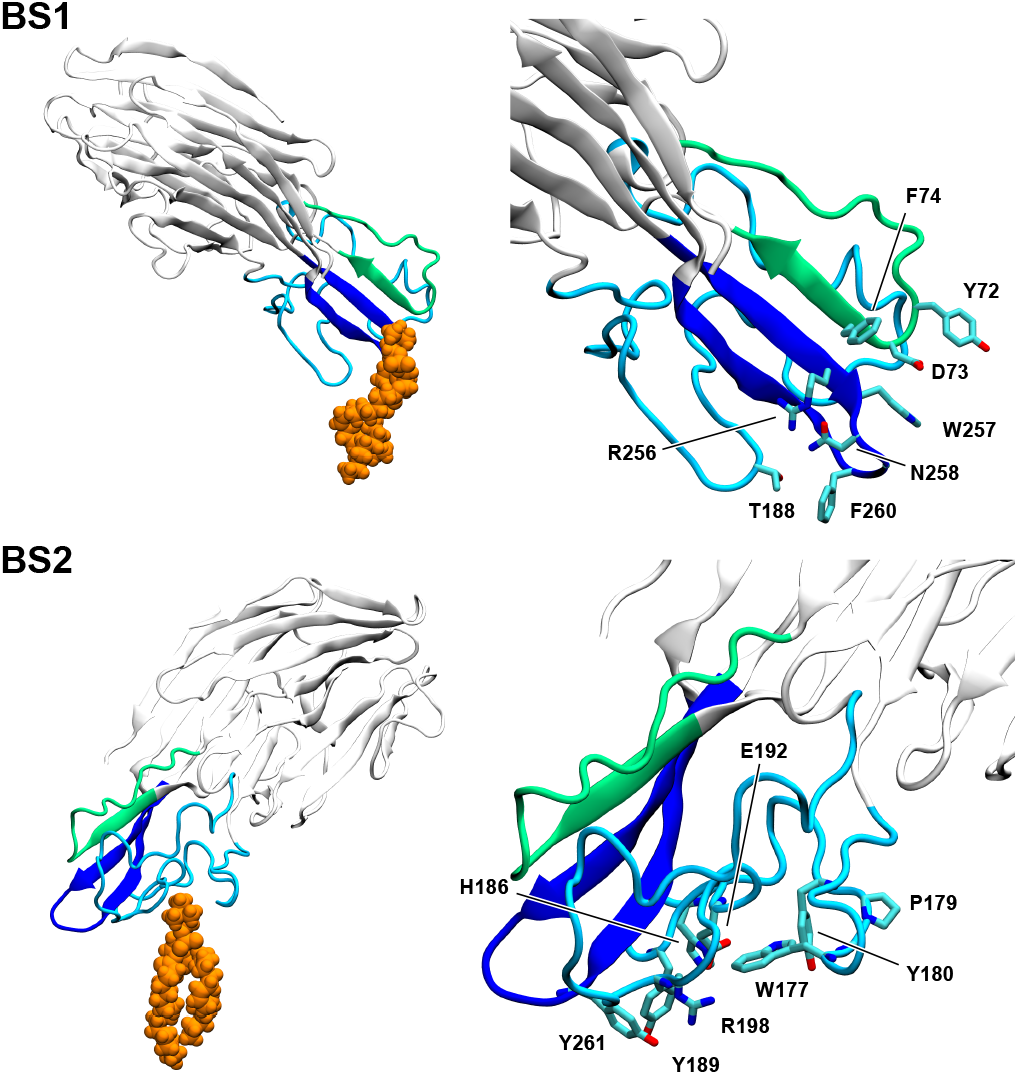
Main DOPC binding sites observed in the simulations of the LukF monomer. The rim domain is colored according to the division in sequences defined in Figure 5, while the bound DOPC molecule, shown in a representative pose, is depicted in orange.

The result from the analysis of the protein-lipids interactions helps rationalize experimental data previously reported in literature. Indeed, a number of identified residues (namely Y72, W177, R198, W257, F260, Y261) had been previously found to be fundamental for LukF binding on the membrane of human erythrocytes [52, 12]; however a comprehensive picture of their interactions with the lipids was missing. The only structural information available in literature concerned the interaction of W177 and R198 with phosphocholine, as captured in the crystal structures of the single LukF monomer [16] and within the assembled γ-HL pore [13]; however, these structures could not explain the role of the other experimentally identified residues.

The binding site BS2 resembles the one known from the crystal structure, where the role of W177 and R198 is preminent in stabilizing the bound PC molecule through cation-π and electrostatic interactions, respectively. BS1, on the other hand, was not captured by previous experimental structures of LukF. In addition, it does not show similarities with the second binding site recently observed in structures of LukD and LukF-PV monomers in complex with C? ?PC. However, the results from our simulations can explain the experimentally observed relevance of residues F260 and Y261 for the membrane binding step of the toxin, while the binding sites emerging from the crystallographic studies of the homologous F components (specifically, LukD and LukF-PV) do not involve such residues [14]. This can be attributed to the intrinsically different binding modes of LukF with respect to LukD and LukF-PV.

## 4. Discussion and conclusions

γ-HL, one of the most common toxins produced by *Staphylococcus aureus,* is composed of two different proteins, LukF and Hlg2. Previous experimental data shed light on the cytotoxic mechanism of action of γ-HL, which consists in the assembly of four LukF and four Hlg2 monomers into an octameric oligomer on the membrane of the target cell, and in the subsequent formation of a transmembrane pore [13, 53]. However, detailed information on the individual steps of pore formation is still missing. Here, we focused on the very first step, namely the binding of γ-HL monomers on lipid bilayers. By collecting and analyzing more than 1.8 ms of CG molecular dynamics simulations, we could observe how the monomer adsorption on the membrane is affected by the exposure of the PC polar heads; the latter, in turn, is modulated by the presence of cholesterol and by the degree of unsaturation of the lipid tails, as shown in atomistic and CG simulations. The calculated dependence of protein absorption on the bilayer composition is in agreement with the experimental data of γ-HL pore formation in model lipid bilayers [10], suggesting that the different binding propensities affecting the first step of pore formation are indeed contributing to the reported experimental observations. In addition, LukF exhibited a larger affinity for the PC-containing bilayer with respect to the Hlg2 component.

More in detail, the collected simulations highlighted the specific role of limited regions of the rim domain in the anchoring process, with some protein residues driving the interaction with the lipids through electrostatic and cation-π interactions. The position of such residues on the protein surface is responsible for the consequent adaptation of preferential orientations of the monomer axis on the membrane bilayer. We remark that the CG model hereby employed adopts the “elnedyn” description for the protein to maintain its secondary structure within the entire simulation; as a consequence, conformational changes of the monomer that might accompany the lipid docking process (such as the possible bending of the cap domain with respect to the rim, or a variation of the conformation of the rim loops) cannot be captured by our CG simulations. Although in principle we cannot rule out the occurrence of such conformational changes, the high similarity of the rim domain in the crystal structures of the monomers and of the protomers in the assembled pore (the RMSD computed on the C_α_ atoms of the rim is approximately 0.3 Å for LukF and 1.3 Å for Hlg2) suggests the absence of large-scale changes in the rim region upon binding. Striking similarities in the structures of the monomer and the protomer were observed also for other leukocidins, as in the case of LukF-PV binding C_14_PC [14].

CG simulations of the monomers in the membrane-bound state allowed us to define the residues of LukF forming the lipid binding site. Remarkably, we found, among them, those that were experimentally identified as relevant for membrane anchoring [52, 12]. In this regard, our study not only corroborates previous experimental results, but also allowed us to rationalize the sparse mutagenesis data in a coherent picture, where the membrane anchoring of the F component is mediated by two distinct binding sites on the rim domain, as observed for the F components of other leukocidins [14]. However, we stress that the identified residues, despite being those driving the first contact between the protein and the membrane, are not necessarily those ensuring a stable anchoring during the oligomerization/perforation phases; the different orientations of the single monomer adsorbed on the membrane and the protomer in the assembled pore suggest indeed an evolution of the contacts between the protein and the bilayer. Further studies are therefore required to assess the role of membrane lipids in the successive phases of pore formation.

### CRediT authorship contribution statement

Conceptualization: TT, GL, RP. Methodology, data collection and analysis: TT. Writing—original draft preparation: TT. Writing—review and editing: TT, GL, RP. Supervision: GL, RP. Funding acquisition: TT, RP. All authors have read and agreed to the published version of the manuscript.

### Data availability

The raw data produced and analysed in this work are freely available on the Zenodo repository at https://doi.org/10.5281/zenodo.6007973.

## Acknowledgements

The authors thank Mauro Dalla Serra for a critical and insightful reading of the manuscript. This project has received funding from the European Research Council (ERC) under the European Union’s Horizon 2020 research and innovation programme (grant agreement No 758588). TT gratefully acknowledges the financial support from the University of Trento through the Seal of Excellence grant.

