## Supplementary information for "Membrane binding of pore-forming γ-hemolysin components studied at different lipid compositions"

### Supporting Information for: Membrane binding of pore-forming $\gamma$-hemolysin components studied at different lipid compositions

Thomas Tarenzi[1,2], Gianluca Lattanzi[1,2], Raffaello Potestio[1,2]

[1] *Physics Department, University of Trento, Via Sommarive 14, Povo (TN), Italy*

[2] *INFN-TIFPA, Trento Institute for Fundamental Physics and Applications, Via Sommarive 14, Povo (TN), Italy*

#### Supplementary figures

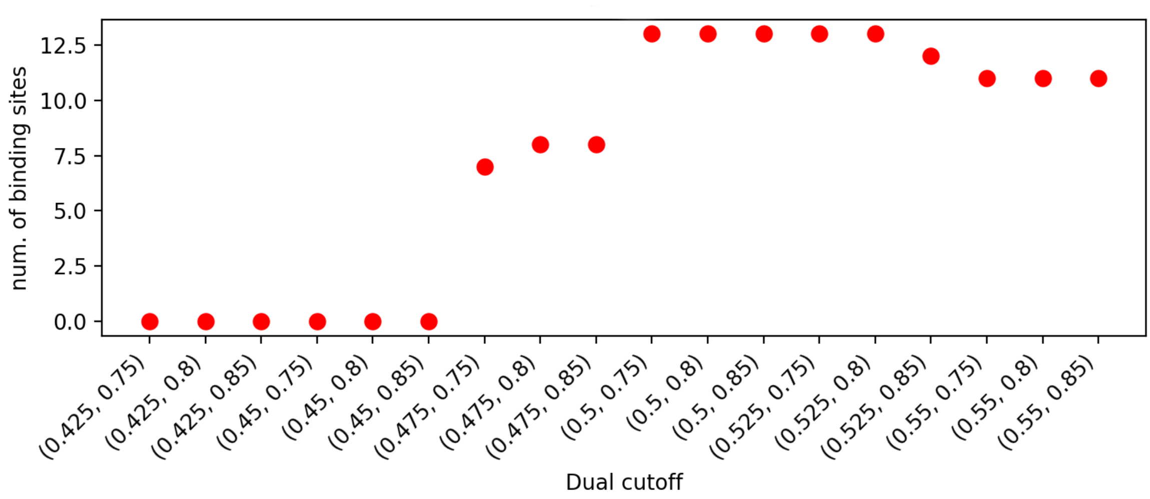


Figure S1: Number of binding sites for DOPC identified form PyLipID for different cutoff values, for LukF docked to DOPC:Chol membrane.

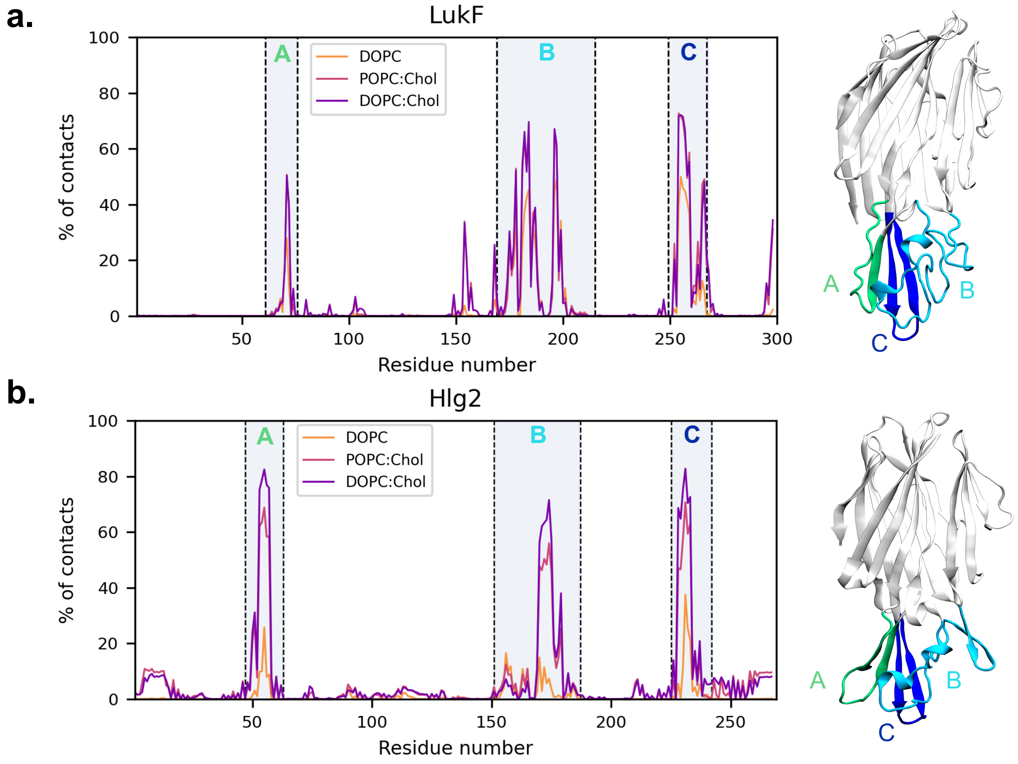


Figure S2: Percentage of frames in which each residue is involved in a contact with the membrane, for the full sequence of LukF (**a**) and Hlg2 (**b**). The three distinct sequences forming the rim domain are also shown on the protein structures for clarity.

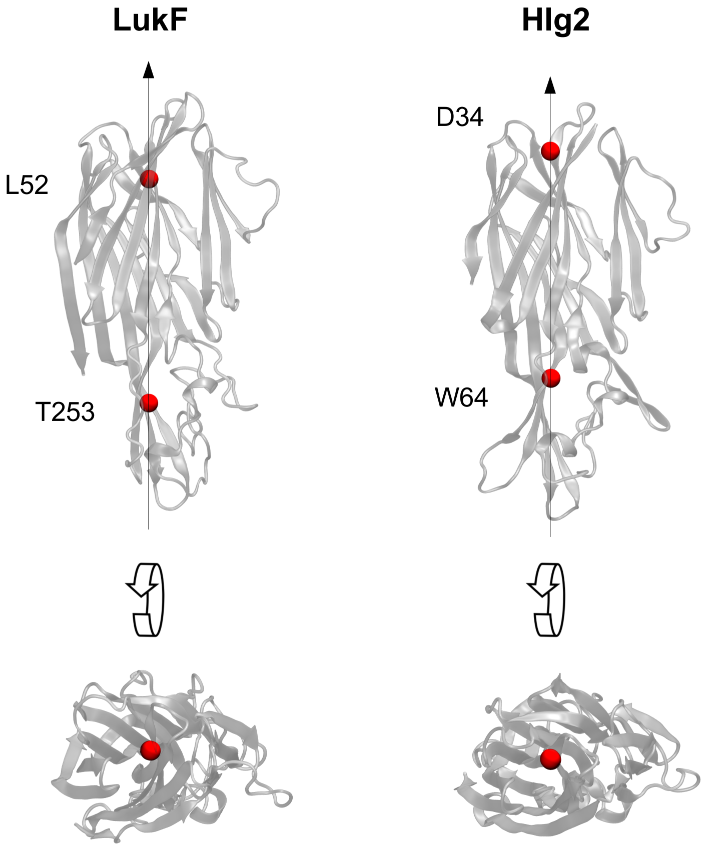


Figure S3: The protein axis used to measure the angle with respect to the normal to the membrane is defined as the axis passing through the backbone beads of residues T253 and L52 for LukF, and residues W64 and D34 for Hlg2.

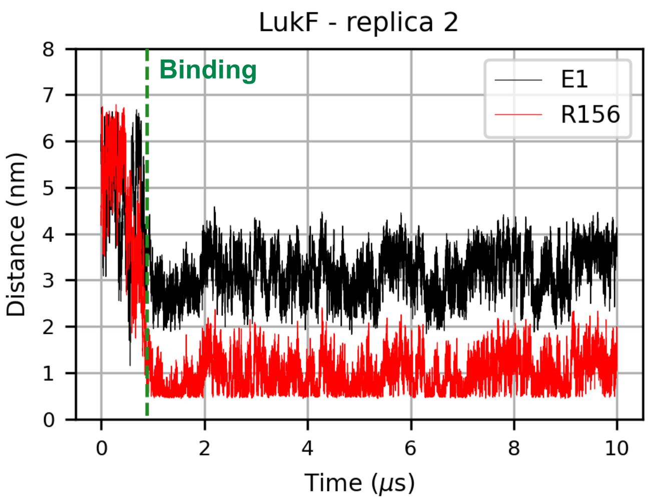


Figure S4: Minimum distance between the bilayer and the backbone beads of E1 (side I) and of R156 (side II) of LukF in Replica 2 of the simulation performed with MARTINI 2.3P force field.

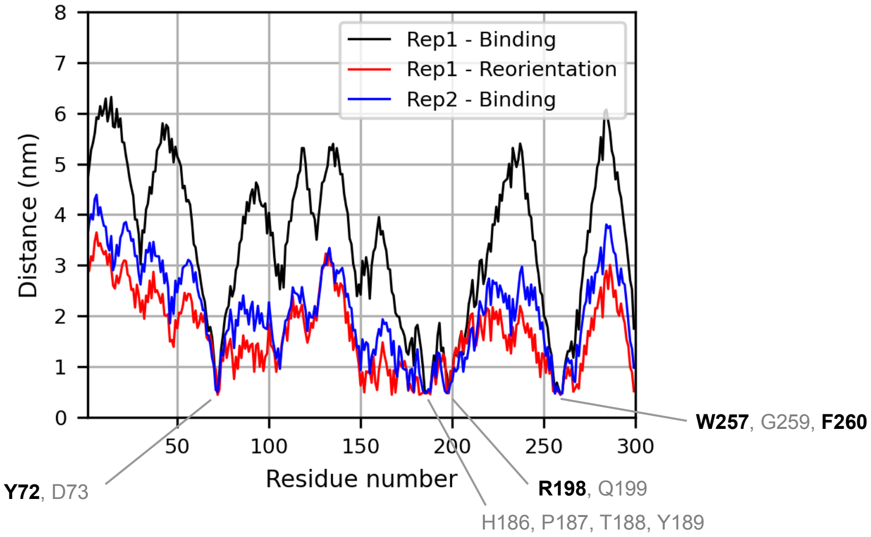


Figure S5: Minimum distance between each residue of LukF and the membrane, in the two replicas performed with MARTINI 2.3P force field. In replica 1, two situations at different times are described (binding=0.85 $\mu$s and reorientation=1.35 $\mu$s of simulation time). The residues that determine the first contacts between the protein and the membrane in both replicas are labelled, and those experimentally found to reduce binding if mutated are in bold [1]. From here, we can also see that the residues driving the first interaction of the toxin with the membrane are indeed located on the rim domain, and include both aromatic and polar/charged residues.

#### Supplementary tables

Table S1: Recap of the atomistic (AT) and coarse-grained (CG) simulations performed in this work. The CG force-field employed is MARTINI 2.2, unless specified otherwise. In all simulations containing the protein and the membrane, the former is placed at a distance of about 1.5 nm from the bilayer surface at the beginning of the simulation.

| System | Resolution | Duration | Number of replicas |
|---|---|---|---|
| DOPC | AT | 1 $\mu$s | 1 |
| POPC:Chol | AT | 1 $\mu$s | 1 |
| DOPC:Chol | AT | 1 $\mu$s | 1 |
| DOPC | CG | 1 $\mu$s | 1 |
| POPC:Chol | CG | 1 $\mu$s | 1 |
| DOPC:Chol | CG | 1 $\mu$s | 1 |
| LukF + DOPC | CG | 3 $\mu$s | 100 |
| LukF + POPC:Chol | CG | 3 $\mu$s | 100 |
| LukF + DOPC:Chol | CG | 3 $\mu$s | 100 |
| Hlg2 + DOPC | CG | 3 $\mu$s | 100 |
| Hlg2 + POPC:Chol | CG | 3 $\mu$s | 100 |
| Hlg2 + DOPC:Chol | CG | 3 $\mu$s | 100 |
| LukF + DOPC:Chol | CG (Martini 2.3P) | 10 $\mu$s | 2 |
| Hlg2 + DOPC:Chol | CG (Martini 2.3P) | 10 $\mu$s | 2 |

Table S2: Comparison between values of surface area per lipid (SA/lipid) as computed from atomistic (AA) and coarse-grained (CG) simulations of the hydrated bilayers, and as measured from previous experiments.

| Composition | AA SA/lipid [$nm^2$] | CG SA/lipid [$nm^2$] | Experimental SA/lipid [$nm^2$] |
|---|---|---|---|
| DOPC | $0.68 \pm 0.01$ | $0.68 \pm 0.01$ | $0.674 \pm 0.005$ [2] |
| | | | $0.676 \pm 0.004$ [3] |
| | | | $0.615 \pm 0.003$ [3] |
| POPC:Chol | $0.42 \pm 0.01$ | $0.43 \pm 0.03$ | $0.451 \pm 0.009$ [4] |
| DOPC:Chol | $0.45 \pm 0.01$ | $0.45 \pm 0.01$ | $0.454 \pm 0.001$ [3] |
| | | | $0.488 \pm 0.004$ [3] |
